# Evaluation of sex differences in laser-induced choroidal neovascularization severity, phenotype and disease progression

**DOI:** 10.1101/2022.04.20.488904

**Authors:** Elizabeth A. Pearsall

## Abstract

Age-related macular degeneration (AMD) progresses through two phases: a non-exudative “dry” stage that compromises vision but does not involve angiogenesis, and an angiogenic “wet” phase characterized by choroidal neovascularization (CNV), in which neovessels from the choroid break through the Bruch’s membrane to form neovascular lesions in the macula. If left untreated, CNV leads to permanent central blindness, which the current standard of care (anti-VEGF therapy) arrests but does not reverse. In laser-induced CNV, a laser is used to rupture the Bruch’s membrane, prompting the outgrowth of choroidal blood vessels in a wound healing response. This model was used to develop currently available CNV therapeutics, such as intravitreal injection of anti-VEGF and corticosteroid therapies and is considered a gold standard model for CNV. Laser-induced CNV thus remains a highly clinically relevant model of CNV secondary to AMD and other pathologies, such as myopia. A rapidly growing body of research suggests that the pathogenesis of myriad diseases is highly dependent on biological sex, yet most studies use only male animals. This highlights the importance of determining whether the severity and pathogenesis of commonly used models of disease, for example laser-induced CNV, differ between sexes. In the present study, we tracked longitudinal progression and pathogenesis of laser-induced CNV in male and female animals, and found that CNV severity did not differ between sexes. This lack of difference recapitulates findings in human patients with CNV, which have also identified that both sexes are at equal risk of CNV development in the contexts of both age-related macular degeneration induced CNV and myopic CNV.

## INTRODUCTION

Aging-related macular degeneration (AMD) is a leading cause of blindness in the elderly population. AMD has two phases, the early non-exudative (dry) phase, characterized by retinal pigment epithelium (RPE) atrophy and cell death in the neural retina [1,2]. In severe cases, dry AMD is followed by wet (exudative) AMD, which is characterized by choroidal neovascularization (CNV) [3]. In CNV, pathological neovessels originating in the choroid plexus invade the macula, a process that leads to irreversible central blindness if left untreated [3,4].

An accumulating amount of evidence suggests that the sequalae of many diseases are highly dependent on male or female sex [5]. Despite robust evidence for sex-specific disease processes, basic and translational research to identify approaches to treatment has historically not taken sex-specific disease pathologies into account [6]. In most studies male animals are used, in part for the sake of convenience, as more female animals are needed to maintain the breeding colony, and excess males can then be used as experimental animals. Women are also underrepresented in many clinical trials, including those that lead to FDA drug approval [7]. Thus, basic mechanistic research and clinical management of disease alike are strongly biased towards male patients; the standard of care for most diseases is based predominately on findings in male animals and patients.

The “NIH Strategic Goals for Women’s Health outlines the need to integrate sex/gender perspectives in basic science and translational research [8] to identify sex-specific disease mechanisms, which can be utilized to develop sex-specific approaches to treatment or potentially to uncover novel disease mechanisms in the context of basic research. Importantly, women are at increased risk for developing major blinding disorders, including AMD [6,9,10]. Women are also more likely to advance from the early “dry” stage of AMD to advanced “wet” neovascular AMD, which is characterized by CNV [11,12]. However, once wet AMD develops, the severity of CNV does not vary between sexes [13,14], nor does the response to anti-VEGF therapy [15,16]. These distinct disease phase-dependent sexual dimorphisms underscore the need to investigate the role of sex in each pathological event of AMD. This could both determine whether sex-specific treatment is necessary for the wet or dry phases of AMD and delineate new disease mechanisms that could potentially be exploited as novel therapeutic approaches.

To meet these goals, it is necessary to determine if sex differences are present in commonly used animal models of both wet and dry AMD, and to report the presence (or absence) of sex-specific differences in disease severity or pathogenesis. No presently available animal models fully recapitulate the process of AMD in humans, particularly the sight threatening CNV characteristic of wet AMD [17,18]. The laser-induced CNV model, in which a laser photocoagulation is used to puncture the Bruch’s membrane and induce pathological CNV in a wound-healing response, is considered a gold-standard model for CNV secondary to wet AMD [17,19]. Laser-induced CNV has been utilized to develop the standard of care for CNV in humans, intravitreal injection of anti-VEGF antibodies, supporting its relevance to human disease [20,21]. In fact, the murine laser-induced CNV model is considered a standard pre-requisite for wet AMD clinical trials [22,23]. Laser-induced CNV has also been used to develop other clinically relevant therapeutic modalities, such as intravitreal injection of corticosteroids [24,25], which is particularly effective in treatment of VEGF non-responders [26,27].

Laser-induced CNV is a progressive model, proceeding through an initial angiogenic phase, which begins by day 3 post-laser photocoagulation, and peaks at day 7 post-photocoagulation [23]. Neovascular lesions begin to regress after day 7, and by day 28 post-laser photocoagulation, resolve nearly completely [22,23]. Contrastingly, in human wet AMD patients, AMD lesions remain stable over time, gradually growing if left untreated, and remaining static if treated with anti-VEGF therapy [2].

In the present study, we evaluated the disease severity and progression of laser-induced CNV in male and female animals. We evaluated multiple disease stages, spanning the initial angiogenic response, the peaks of lesion edema and vascular leakage, the peak of neovessel formation, and subsequent regression of CNV lesions. Commonly used CNV outcomes, such as optical coherence tomography (OCT)-measured lesion volume, fluorescein angiography (FA) assessment of lesion leakage, and histological quantification of lesion size were assessed in both sexes and across time points.

Determining the presence (or absence) of a sex-specific response to laser-induced CNV is important for elucidating basic disease mechanisms. Further, because the laser-induced CNV model is commonly used to develop AMD therapeutics for clinical use, it is essential that this model be standardized across investigators, and that potential sources of biological variation be rigorously evaluated and reported to the research community. This study is therefore of significant relevance not only to basic research, but also to preclinical research aimed at developing novel wet AMD therapeutics.

## MATERIALS AND METHODS

### Experimental animals

Seven-week-old male and female C57 bl6/J mice were purchased from Jackson Labs (Bar Harbor, Maine). Mice were maintained in specific pathogen-free conditions and given standard laboratory diet and tap water ad libitum. After allowing 1 week for acclimation, mice were subjected to laser photocoagulation induction of CNV, and to imaging analyses as described below, before euthanasia on either day 7 or day 14 post-CNV induction. All animal handling and experimentation were pre-approved by the Massachusetts Eye & Ear Institutional Animal Care and Use Committee prior to beginning experiments (approval no. 14-067) and were conducted in accordance with the NIH Guidelines for the Care and Treatment of Laboratory Animals and the ARVO Statement for the Use of Animals in Ophthalmic and Vision Research. Animals were euthanized with anesthesia overdose (Avertin, Sigma, St. Louis, MO, 200 mg/kg) and subsequent cervical dislocation after deep anesthesia was confirmed. This euthanasia protocol was consistent with the recommendations of the AVMA guidelines for the euthanasia of animals.

### Experimental numbers and statistical analyses

Experiments were conducted in groups of n = 5 mice/group. Experiments were performed at minimum in duplicate (day 14 time point) and at maximum in quadruplicate (days 3–7 time points). Therefore, the minimum number of animals was at least n = 10 mice/group for all treatment groups and time points. For pairwise comparisons of numerical values, significance was assessed with an unpaired, two-tailed Student’s t-test, and p < 0.05 was considered statistically significant. For multiple comparisons of numerical values, a one-way ANOVA with Tukey’s posthoc comparison was used. To assess the significance of categorical variables between groups, a chi-squared (χ^2^) test was used. For statistical analyses, “n” is reflective of individual lesions unless otherwise specified. Data are expressed as means ± standard error of the means (S.E.M.). Statistical outliers were identified using the ROUT test, CI 90%, and were excluded from final analyses.

### Laser photocoagulation and CNV induction

CNV was induced as described previously [28]. Briefly, a 532 nm laser (Oculight GLx Laser System, IRIDEX) attached to a slit lamp was used for photocoagulation. In order to view the posterior pole of the eye, a coverslip with a drop of Goniosol was applied to applanate the cornea. For size and leakage studies, four laser-induced lesions are made around the optic nerve in a clock-wise manner (3, 6, 9, and 12 o’clock). The laser was set as follows: 100 mW power, 100 μm spot size, and 0.1s pulse duration. The appearance of a bubble at the site of photocoagulation was used to identify Bruch’s membrane disruption, and therefore any laser injury without a corresponding bubble was excluded from analysis. Mice underwent live-animal imaging outlined below on days, 3, 5, 7, and 14 after laser photocoagulation/CNV induction to quantify lesion size and leakage in live animals. At 7 or 14 days post-laser photocoagulation/CNV induction, mice were euthanized and eyes were collected for choroidal flat-mount.

### Fluorescence angiography and lesion leakage assessment

A Micron IV Retinal Imaging Microscope (Phoenix Research Laboratories, Pleasanton, CA) was used to capture fluorescein images as described previously [28]. First, mice were anesthetized with Avertin (200 mg/kg, Sigma, St. Louis, MO) and kept on a heating pad (37°C) to maintain body temperature. 5% phenylephrine and 0.5% tropicamide were used for pupil dilation. Animals were injected intraperitoneally (ip) with 0.1 ml 2% fluorescein sodium (Akorn, Lake Forest, Il). The corneas were covered with 2.5% Goniovisc (HUB Pharmaceuticals, Rancho Cucamonga, CA) and subsequently placed in contact with the Micron camera lens. Images were obtained 3-5 minutes (early phase) and 7-10 minutes (late-phase) after Fluorescein injection using StreamPix software (Phoenix Research Laboratories, Pleasanton, CA). A previously established scheme was used to grade the hyperfluorescent lesions: faint or mottled fluorescence without leakage was scored as “0” (no leakage); hyperfluorescence without any increase in size or intensity between early- and late-phase images was scored as “1” (debatable leakage); hyperfluorescence with consistent size but increasing intensity was scored as “2A” (definite leakage); and hyperfluorescence with increasing size and intensity was scored as “2B” (clinically significant leakage) (Fig. S1) [28].

### Optical coherence tomography (OCT)

A Bioptigen system was used to conduct spectral domain-optical coherence tomography (SD-OCT) as described previously [29]. Briefly, mice were first anesthetized and placed on a freely rotating cassette customized for optimal eye alignment. AB-scans with an area size of 1.4 × 1.4 mm were obtained using 100 horizontal, raster, and consecutive B-scan lines, each composed of 1,000 A-scans. Cross-sectional images of the lesions were captured at the consecutive B-scan line with the largest lesion size, and lesion volume of cross-sections was then measured using NIH Image J software.

### Choroidal Flatmounts

Choroidal flatmounts and measurement of lesion size were conducted as previously [28]. At 7 or 14 days post-photocoagulation, mouse eyes were enucleated and fixed in 4% paraformaldehyde. Whole retinas were then separated from the underlying choroid and sclera (eyecup). BSA/Triton X (2% BSA, 0.1% Tx) was utilized to permeate the lesions prior to staining (5 hours at room temperature or overnight at 4°C). Alexa Fluor 488 conjugated to isolectin B4 (1:100, Invitrogen) was used to stain the eyecups overnight at 4°C. PermaFluor aqueous mount (Thermo Scientific) was then used to flat-mount eyecups with the choroid facing upward and the sclera facing downward. Flat-mount images were captured using a Zeiss AxioCam MRm camera and Zeiss AxioObserver (Z1) microscope, and ImageJ software (National Institutes of Health) was used to quantify lesion size.

### Exclusion Criteria

All images will be blinded prior to analysis. The following pre-determined exclusion criteria were applied after the images were blinded. For flat-mount images, overburnt lesions and fused lesions were excluded. For OCT, lesions with subretinal edema or greater than 20 μm in size were excluded. For FA, fused lesions were excluded.

## RESULTS

CNV lesion volume and edema (OCT) and lesion leakage (FA) were measured in live animals across multiple phases of laser-induced CNV disease progression and regression (days 3-, 5-, 7-, and 14-post laser photocoagulation). CNV lesion size was measured using histology during later phases of CNV (days 7 and 14 post-CNV induction).

### Early angiogenic response (D3)

In the murine laser-induced CNV model, the angiogenic response begins by day 3 post-photocoagulation (D3), with VEGF production beginning on D2. We thus quantified lesion volume and leakage beginning on D3. Lesion leakage did not vary between sexes (Fig. 1A-C, χ^2^ p > 0.05), nor did lesion volume (Fig. 1D-E, p > 0.05, Student’s t-test).

**Figure 1:**
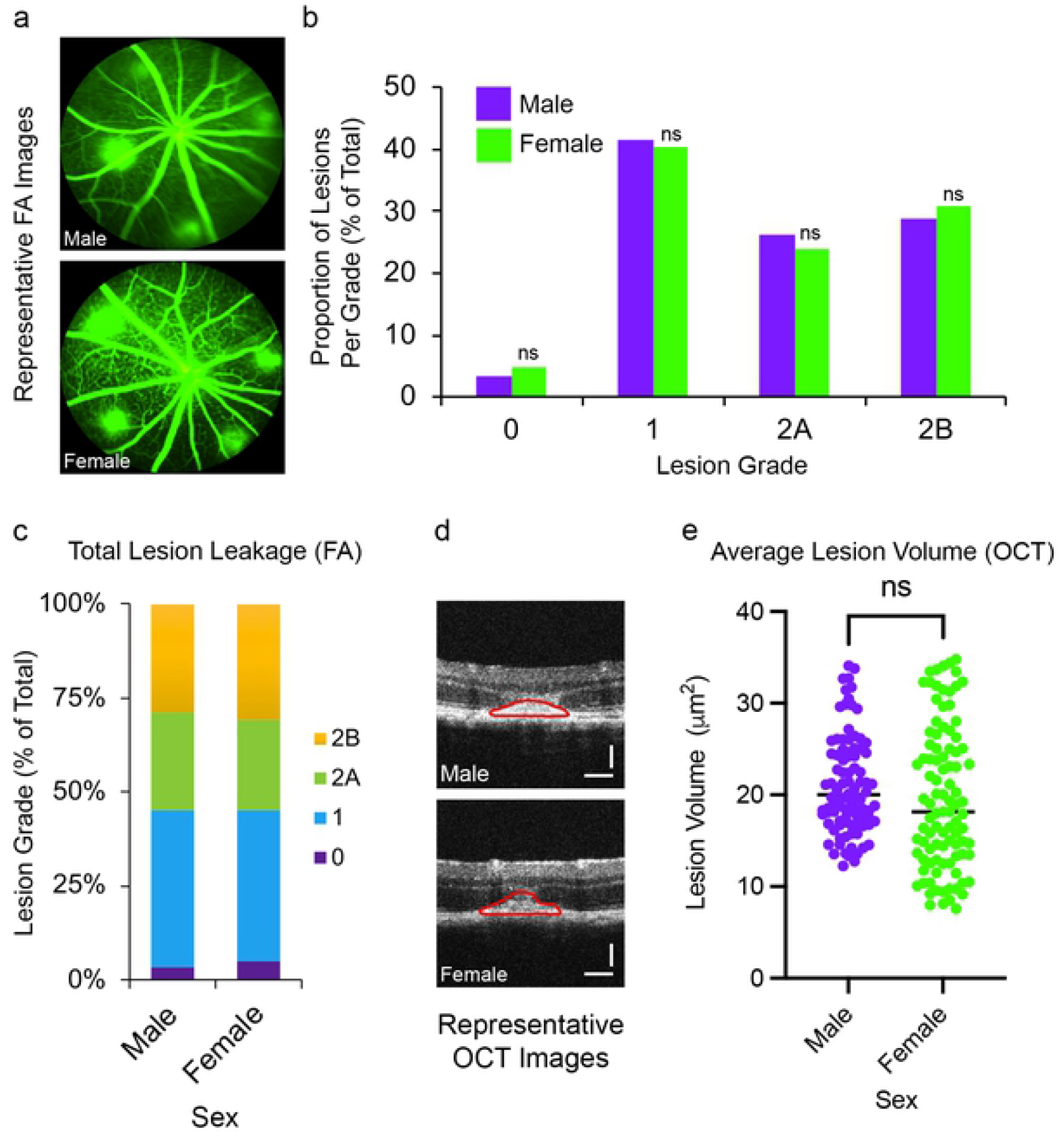
Similar early angiogenic response following laser-induced choroidal neovascularization (CNV) induction in male and female animals. **(a)** Representative fluorescein angiography (FA) images of male and female animals on day 3 post-CNV. **(b-c)** Relative proportions of FA lesion grades were similar in male and female animals on day 3 post-CNV induction (male n=149, female n=146 lesions, χ^2^ p ≥ 0.05, not significant (ns)). **(d)** Representative optical coherence tomography (OCT) images on day 3 post-CNV. **(e)** Lesion volume, as measured by OCT, did not differ significantly in male and female animals on day 3 post-CNV induction (male n=93, female n=97 lesions, one-sided two-tailed Student’s t-test p ≥ 0.5, not significant (ns)).

**Figure 2:**
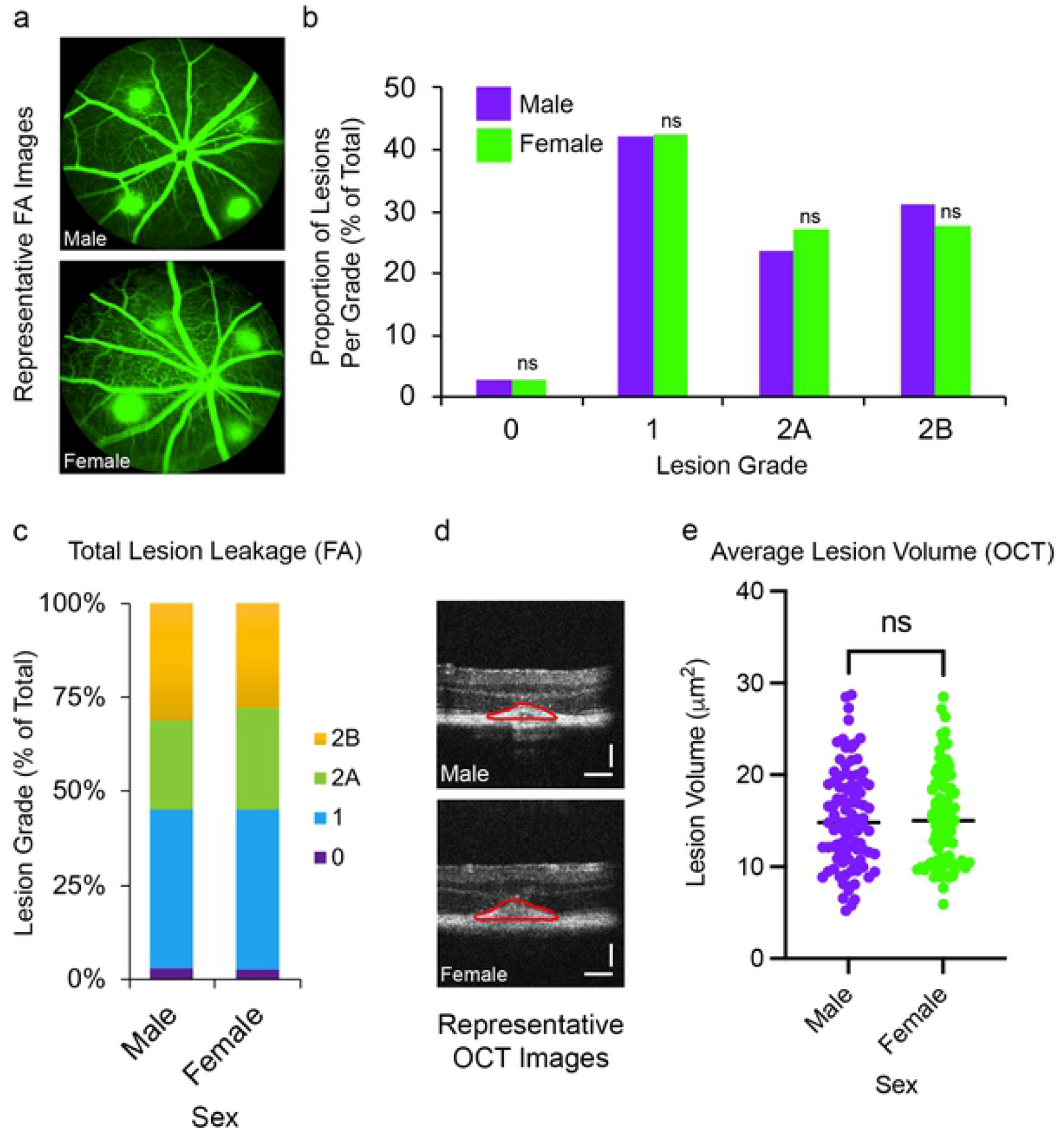
Similar mid-stage CNV lesion growth and leakage in male and female animals. **(a)** Representative FA images of male and female animals on day 5 post-CNV. **(b-c)** Relative proportions of FA lesion grades were similar in male and female animals on day 5 post-CNV induction (male n=135, female n=144 lesions, χ^2^ p ≥ 0.05, ns). **(d)** Representative optical coherence tomography (OCT) images on day 5 post-CNV. **(e)** Lesion volume, as measured by OCT, did not differ significantly in male and female animals on day 5 post-CNV induction (male n=94, female n=84 lesions, one-sided two-tailed Student’s t-test p ≥ 0.5, ns).

### Mid-stage disease (D5)

On D5 post-CNV, neither lesion leakage (Fig. 3A-C, χ^2^ p > 0.05) nor lesion volume (Fig. 3D-E, Student’s t-test p < 0.05) differed between sexes. This suggested that even when lesion leakage and edema were maximal, there was no significant difference in these parameters between sexes.

**Figure 3:**
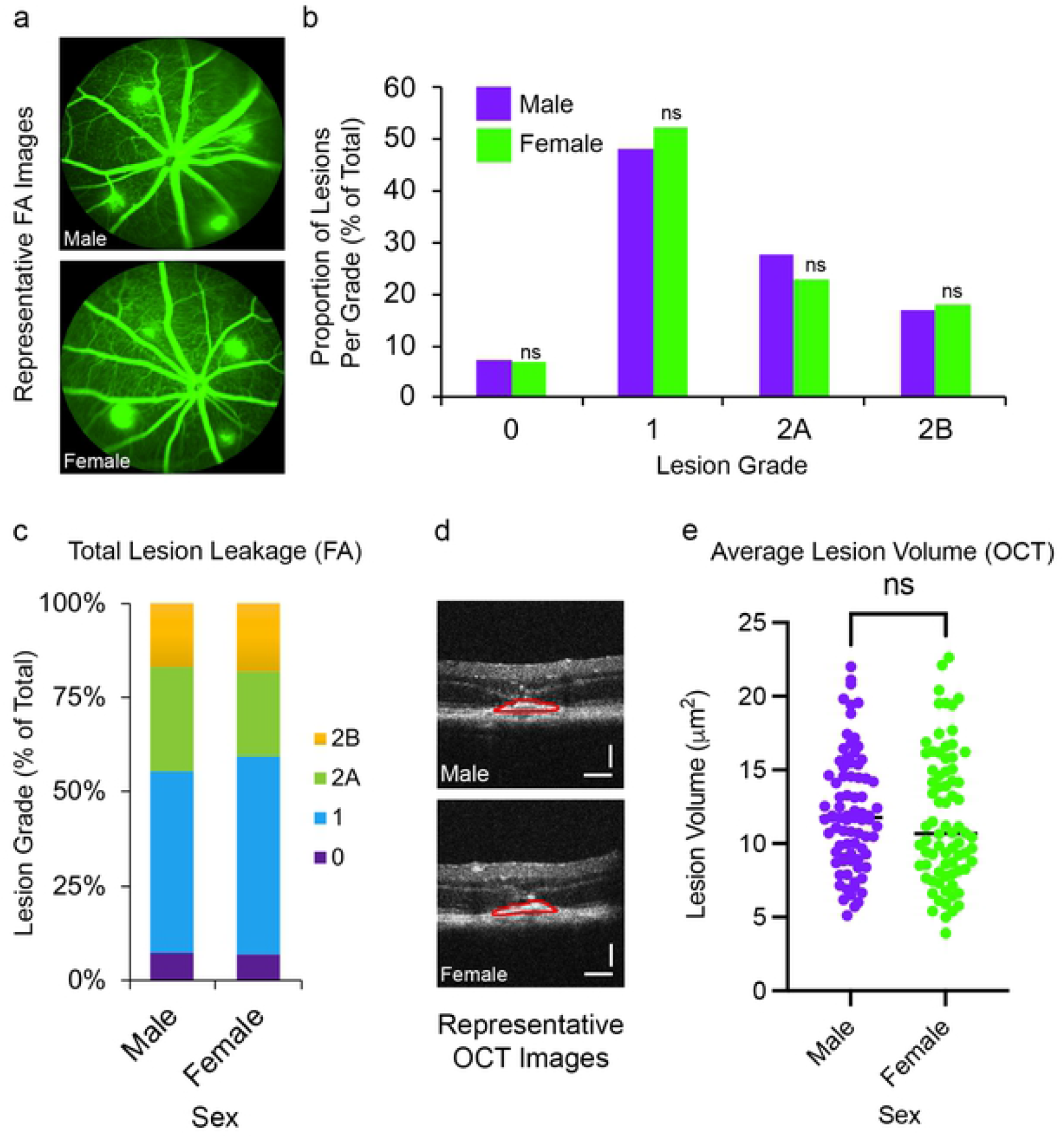
Similar peak angiogenic response following CNV induction in male and female animals. **(a)** Representative FA images of male and female animals on day 7 post-CNV. **(b-c)** Relative proportions of FA lesion grades were similar in male and female animals on day 7 post-CNV induction (male n=137, female n=145 lesions, χ^2^ p ≥ 0.05, ns). **(d)** Representative optical coherence tomography (OCT) images on day 7 post-CNV. **(e)** Lesion volume, as measured by OCT, did not differ significantly in male and female animals on day 7 post-CNV induction (male n=80, female n=76 lesions, one-sided two-tailed Student’s t-test p ≥ 0.5, ns).

### Peak neovascularization (D7)

Histological assessment of CNV lesions labels blood vessels to exclusively evaluate outgrowth of pathological neovessels. Prior studies have demonstrated that this metric is highest at D7 post-CNV [30]. At this time point, neither lesion leakage (Fig. 4A-C, χ^2^ p > 0.05) nor lesion volume (Fig. 4D-E, Student’s t-test p > 0.05). Consistently, histologically measured lesion size did not vary between sexes on D7 (Fig. 6A-B, p > 0.05, Student’s t-test). This suggested that at peak neovascularization, there was no significant difference between sexes.

**Figure 4:**
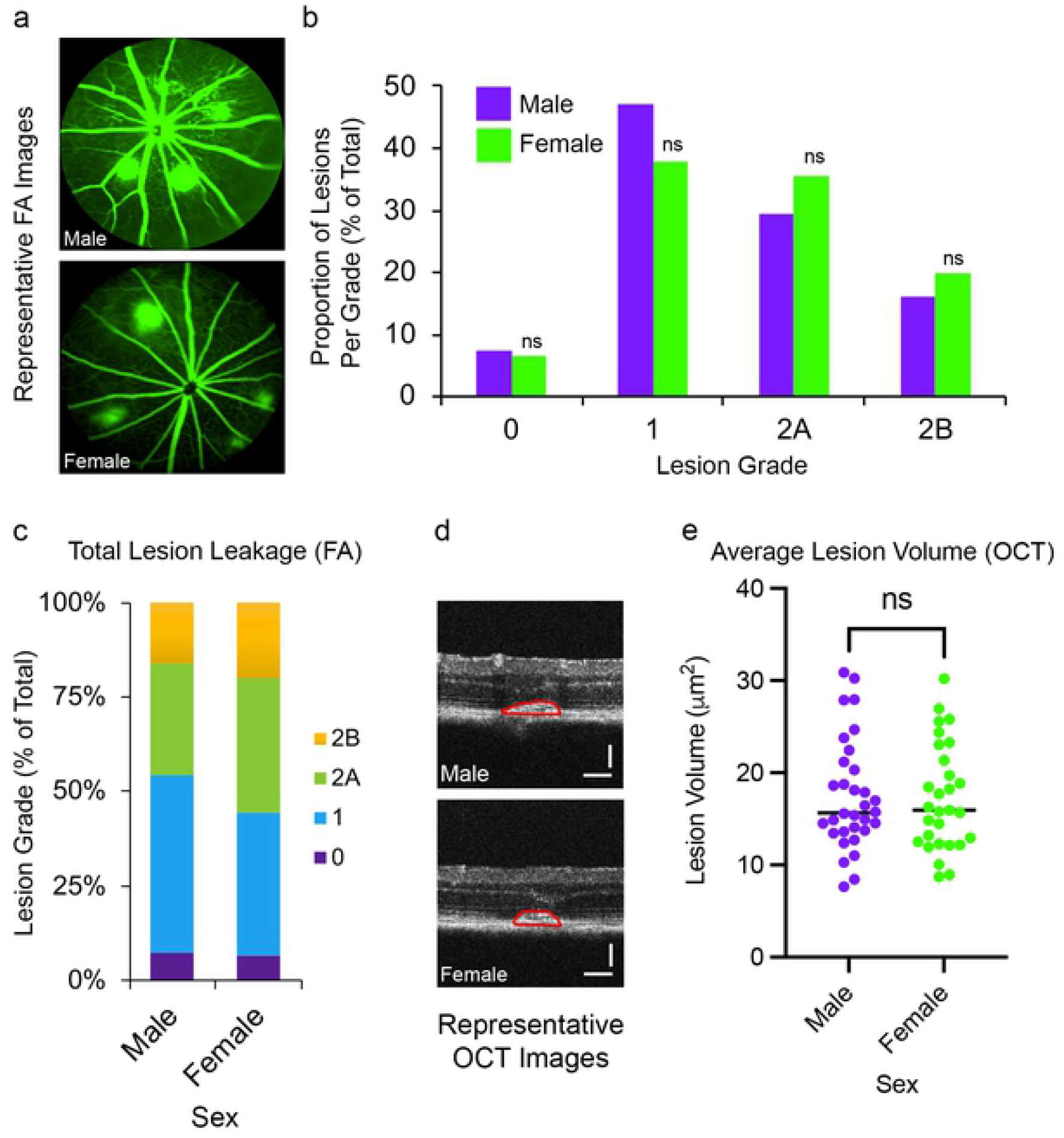
Similar lesion regression following CNV induction in male and female animals. **(a)** Representative FA images of male and female animals on day 14 post-CNV. **(b-c)** Relative proportions of FA lesion grades were similar in male and female animals on day 7 post-CNV induction (male n=68, female n=45 lesions, χ^2^ p ≥ 0.05, ns). **(d)** Representative optical coherence tomography (OCT) images on day 14 post-CNV. **(e)** Lesion volume, as measured by OCT, did not differ significantly in male and female animals on day 14 post-CNV induction (male n=32, female n=29 lesions, one-sided two-tailed Student’s t-test p ≥ 0.5, ns).

### CNV regression (D14)

After laser photocoagulation in mice, CNV begins on approximately D3, and progressively increases, reaching peak NV on D7. After this timepoint, lesion size and pathology begin to decrease in severity [30]. During the regressive phase of CNV (D14), neither lesion leakage (Fig. 5A-C, χ^2^ p > 0.05) nor lesion volume (Fig. CD, p > 0.05, Student’s t-test) differed between sexes. Similarly, lesion size as quantified by histology did not differ between sexes at this timepoint (Figure 6C-D, p > 0.05, Student’s t-test). Collectively, these findings demonstrated that biological sex did not affect CNV regression.

**Figure 5:**
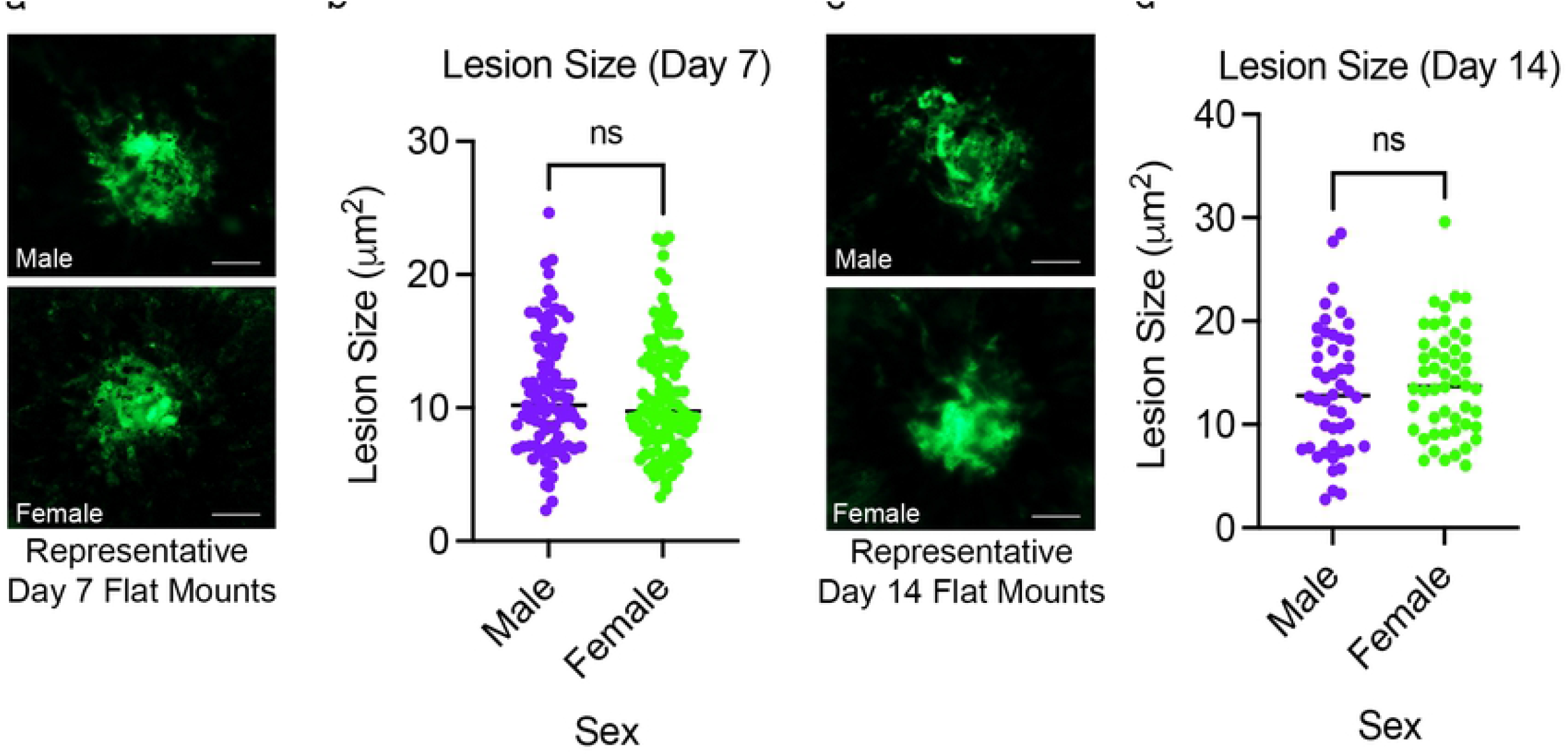
Similar lesion sizes in early and late-stage CNV in male and female animals. **(a)** Representative choroidal flat-mount images on day 7 post-CNV. **(b)** Lesion size, as measured by choroidal flat mount, was similar in male and female animals on day 7 post-CNV (male n=97, female n=111 lesions, one-sided two-tailed Student’s t-test p ≥ 0.5, ns). **(c)** Representative choroidal flat-mount images on day 14 post-CNV. **(d)** Lesion size, as measured by choroidal flat mount, was similar in male and female animals on day 14 post-CNV (male n=48, female n=50 lesions, one-sided two-tailed Student’s t-test p ≥ 0.5, ns).

## DISCUSSION

The basic biology of males and females is vastly different, and it is thus unsurprising that the presentation, pathology, and mechanisms of myriad diseases are sexually dimorphic [5]. For example, sexual dimorphism has been well-documented in cardiovascular and autoimmune disease [31–34], yet most basic research studies use only one sex of animals, and the standard of care and clinical management for most diseases, including AMD, is the same for both sexes. Because basic research uses predominately male animals and women are underrepresented in clinical studies [7], treatment and clinical management of most diseases is biased towards males.

In humans, AMD proceeds through two phases: a non-exudative “dry” phase characterized by atrophy of the retinal pigment epithelium and photoreceptor dysfunction in the macula, which is in some cases followed by exudative “wet” AMD, characterized by pathological neovessel formation in the choroid (CNV) [2]. Although dry AMD compromises visual function parameters such as contrast sensitivity, reading speed, microperimetry and dark adaptation, progression to wet AMD is the primary sight-threatening event in AMD and can lead to permanent central blindness if left untreated. Thus, most currently available approaches to treatment target wet AMD, as these pathologies have the most profound effects on patient quality of life. Preventing or alleviating CNV are also primary targets for ongoing research efforts to identify new therapeutic approaches. Women are more likely to develop AMD, beginning with the onset of dry AMD pathologies such as RPE atrophy, drusen formation and photoreceptor degeneration [6,9,10]. Women are also more likely to progress from dry AMD to wet AMD [11,12,35]. However, once CNV develops, neither the severity of CNV nor the response to anti-VEGF therapy differs between sexes [15,16]. However, because women are underrepresented in clinical and epidemiological studies, and are less likely to seek medical treatment, including for ophthalmic disease [6], epidemiological studies should be interpreted with caution.

Notably, the innate immune response is more robust in females [36], and pathological activation of innate immunity is a known risk factor and contributor to AMD [37]. The adaptive immune response is also more robust in females [38], and some studies suggest that adaptive immunity can also contribute to AMD [39,40]. Women are also more susceptible to most age-related diseases [36], which in some cases has been linked to the radical hormonal changes of menopause.

The disease stage-dependent sexual dimorphism of AMD necessitates separate investigation of each pathological event of wet and dry AMD. Laser-induced CNV is a commonly used model of the CNV that occurs in advanced wet AMD. Although lesions are generated by artificial disease induction, the disease processes by which neovessels invade disruptions in the Bruch’s membrane has significant commonalities with the neovascular events of wet AMD [17,19]. The model’s relevance to human disease is underscored by its use to generate multiple clinically effective interventions [20,21,24,25], and laser-induced CNV is, in fact, a prerequisite for any clinical trials involving anti-VEGF therapies [22,23].

Despite being the gold standard model for wet AMD, laser-induced CNV is subject to significant investigator-dependent variables, which must be minimized and standardized to ensure consistency of the model across research groups. CNV lesions are induced by laser photocoagulation in the laboratory, and experimental outcomes can be affected not only by variations in the photocoagulation process itself, but also by the quantification methods used. Important prior studies have optimized these processes, significantly improving the consistency of disease induction and quantification [30,41]. The progression and severity of laser-induced CNV can also be affected by biological variables such as housing conditions, diet, and sex. For example, other studies have demonstrated that dietary lipids significantly affect retinal and choroidal neovascularization [42–44], necessitating further investigations into this potential variable and the standardization of laboratory diets used in CNV studies, particularly the source of lipids. No prior studies have directly assessed the effect of biological sex on laser-induced CNV, which was the objective of the present study.

We demonstrated that sex did not affect common measures of CNV disease severity, including lesion leakage (angiography), lesion volume (OCT), and lesion size (IHC). Moreover, disease severity did not differ between sexes across multiple stages of CNV development and progression (Days 3–7), nor during disease regression (Day 14). This would be consistent with human data, as epidemiological studies suggest that the severity and treatment response rate of CNV does not differ between male and female wet AMD patients [13–16]. However, the development of AMD and progression to CNV is more common in women [6,9–12], suggesting that potential sex differences should also be examined in animal models of dry AMD and spontaneous CNV. Further, age-dependent sex differences in the severity of laser-induced CNV cannot be ruled out.

Although the present findings are highly significant to standardization of the laser-induced CNV model, they should still be interpreted cautiously, and potential sex differences in specific disease processes or putative therapeutics should still be assessed. For example, microglia contribute significantly to the progression and severity of laser-induced CNV [45–48], and prior studies have suggested that microglial biology significantly differs between sexes [49,50]. Other processes of the inflammatory response to CNV, for example peripheral macrophage and neutrophil-mediated innate immunity [51–53], also differ significantly between sexes in other disease processes [38,54]. Thus, although disease severity does not vary between sexes in the base model of murine laser-induced CNV, future studies of specific disease mechanisms or targeted therapeutics using this model should at minimum rule out the potential for sex differences.

## ACKNOWLEDGEMENTS

This work was supported by EY029962 (EP). The author thanks Drs. Dong Ho Park and Kip M. Connor for their contributions to this work.

